# Osteoblasts pattern endothelium and somatosensory axons during zebrafish caudal fin organogenesis

**DOI:** 10.1101/2021.09.28.462226

**Authors:** Rosalind G. Bump, Camille E. A. Goo, Emma C. Horton, Jeffrey P. Rasmussen

**Affiliations:** Department of Biology, University of Washington, Seattle, WA, 98195, USA; Molecular and Cellular Biology Program, University of Washington, Seattle, WA, 98195, USA; Developmental Biology and Stem Cell Program, University of California, San Francisco, CA 94143, USA

**Keywords:** *Danio rerio*, bone, somatosensory neuron, neurovascular congruence, VEGFR, developmental biology

## Abstract

Skeletal elements frequently associate with vasculature and somatosensory nerves, which regulate bone development and homeostasis. However, the deep, internal location of bones in many vertebrates has limited *in vivo* exploration of the neurovascular-bone relationship. Here, we use the zebrafish caudal fin, an optically accessible organ formed of repeating bony ray skeletal units, to determine the cellular relationship between nerves, bones, and endothelium. In adults, we establish the presence of somatosensory axons running through the inside of the bony fin rays, juxtaposed with osteoblasts on the inner hemiray surface. During development, we show the caudal fin progresses through sequential stages of endothelial plexus formation, bony ray addition, ray innervation, and endothelial remodeling. Surprisingly, the initial stages of fin morphogenesis proceed normally in animals lacking either fin endothelium or somatosensory nerves. Instead, we find that *sp7+* osteoblasts are required for endothelial remodeling and somatosensory axon innervation in the developing fin. Overall, this study demonstrates that the proximal neurovascular-bone relationship in the adult caudal fin is established during fin organogenesis and suggests that ray-associated osteoblasts pattern axons and endothelium.

**Summary statement:** Analysis of cellular interdependence during caudal fin development reveals roles for osteoblasts in patterning endothelium and somatosensory axon innervation.

## Introduction

Vertebrates display a vast diversity of form and function but share a common biological organization—bony structures innervated by dense networks of somatosensory nerves and blood vessels. The neurovascular-bone connection is a long-standing paradigm in vertebrate biology, noted as early as the 19^th^ century (Berthold, 1831). This relationship is observed in diverse species with convergent, innervated features like skeletal long bones (Tomlinson et al., 2016) and teeth (Shadad et al., 2019), as well as divergent, specialized features like deer antlers (Wislocki and Singer, 1946) and elephant tusks (Weissengruber et al., 2005). Sensory nerves detect stimuli critical for survival (e.g., pain) and blood vessels deliver oxygen and essential nutrients to the bones.

Growing evidence indicates that vasculature and peripheral nerves play key roles in bone development and homeostasis. For example, the development of cranial foramina, or passageways of the skull, has been proposed to rely on sensory nerves and vasculature to restrict ossification to discrete domains (Akbareian et al., 2015). Additionally, damage to bony structures results in mis-patterned, incomplete repair when the tissue is denervated, as observed in deer antlers (Suttie and Fennessy, 1985), mouse femurs (Chen et al., 2019), and axolotl teeth (Makanae et al., 2020). In humans with congenital insensitivity to pain, a condition arising from an incomplete peripheral nervous system, patients experience significant musculoskeletal disorders, including repeated fractures (Bar-On et al., 2002; Mifsud et al., 2019). Similarly, inhibition of angiogenesis limits bone repair in several contexts (Hausman et al., 2001; Street et al., 2002). Taken together, these data strongly indicate the necessity of a functional neurovascular unit to promote the morphogenesis and maintenance of osteogenic tissue.

Despite how critical the neurovascular-bone axis is for organismal form and function, the deep, internal localization of bones has presented a barrier to *in vivo* study in mammalian models. By contrast, the superficial bony structures of teleost (ray-finned) fish offer an attractive alternative to overcome the physical barriers to studying this highly conserved relationship. In particular, the teleost caudal fin is a thin, optically accessible organ, composed of a reproducibly patterned set of dermal bones known as fin rays (lepidotrichia) (Becerra et al., 1983; Bird and Mabee, 2003; Montes et al., 1982). In adult zebrafish, the caudal fin rays associate with blood vessels and contain axon-associated Schwann cells (Lee et al., 2013; Xu et al., 2014). However, the identity of the fin nerves and their relationship to other tissues during fin morphogenesis remains poorly understood.

Here, we leverage the imaging and genetic advantages of zebrafish to investigate the establishment of the neurovascular-bone relationship in caudal fin bony rays. We show that somatosensory axons innervate the fin rays in close association with ray osteoblasts. We establish the developmental timing of ray innervation relative to endothelial and osteoblast remodeling during fin morphogenesis. Surprisingly, we report that neither bony rays nor axon development depend on the presence of an endothelial plexus within the caudal fin. Similarly, the initial stages of fin organogenesis proceed normally in the absence of somatosensory nerves. However, we find that conditional ablation of osteoblasts disrupts endothelial and axon morphogenesis. Together, our findings point to a central role of osteoblasts in establishing the precise patterning of endothelium and axons during fin organogenesis.

## Results

### Adult peripheral axons innervate each bony ray of the caudal fin

The adult zebrafish caudal fin has a highly stereotyped and well-described bony anatomy (Fig. 1A). The bi-lobed fin contains 18 main bony rays, with each ray built from individual bone segments. With the exception of the dorsal-most and ventral-most rays, each bony ray bifurcates (branches into two) as the rays extend distally (Fig. 1A,B). The bony rays do not form solid cylinders of calcified bone, but rather comprise two concave “hemirays”. The caudal fin contains a wide range of cell types with diverse developmental origins, both in the interray space and within the hemirays themselves. For example, a monolayer of bone-depositing osteoblasts covers each hemiray; an artery runs through the middle of each ray, while veins flank the dorsal and ventral margins of each ray (Xu et al., 2014); and previous analyses of cell lineages within the caudal fin identified pigment cells, fibroblasts, and neural crest-derived Schwann cells running through the rays (Lee et al., 2013; Tu and Johnson, 2011). Additionally, staining of fins with general axonal markers indicates axons innervate the adult fin (König et al., 2019; Lisse et al., 2016; Lee et al., 2013). However, the identity of these axons and the relationship between nerves and other anatomical structures in the caudal fin remains poorly described.

**Figure 1.**
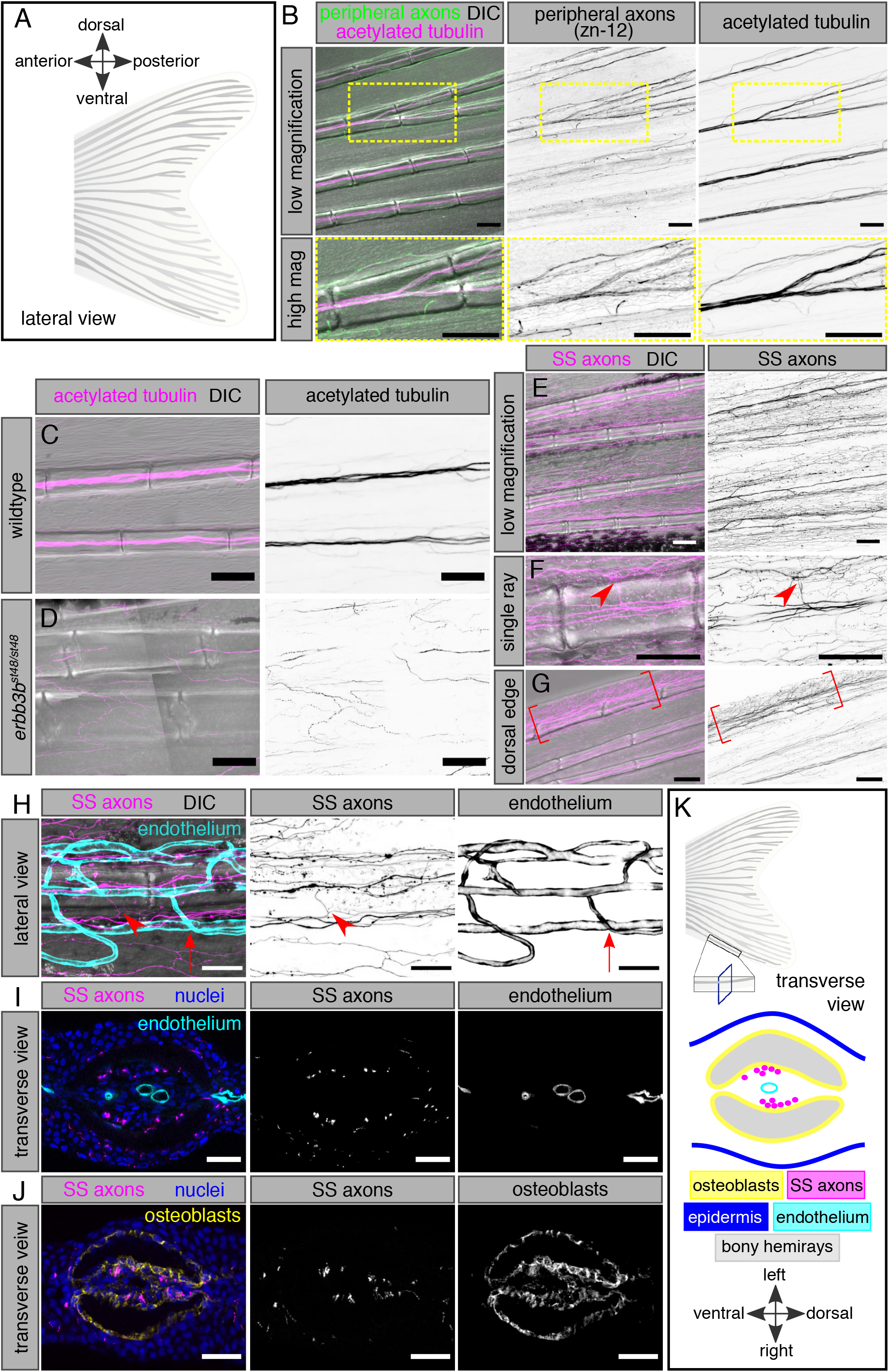
Somatosensory innervation of the adult caudal fin. **(A)** Schematic of the adult zebrafish caudal fin anatomy, composed of 18 bony rays. Posterior (distal) is to the right in this and subsequent lateral views. **(B)** Lateral view of adult caudal fin, fixed and immunostained for peripheral axons (zn-12 antibody) and acetylated tubulin. High magnification insets of a ray bifurcation. Note that zn-12 preferentially stains axon endings, whereas acetylated tubulin preferentially labels axon bundles. **(C**,**D)** Representative images of caudal fin bony rays (DIC) and acetylated tubulin staining in adult fins of the indicated genotypes. **(E-G)** Caudal fin bony rays (DIC) and somatosensory axons (SS axons) of an adult *Tg(p2rx3a>mCherry)* fish. The arrowhead in F illustrates an axon branching to exit the ray segment. Note the increased density of somatosensory axons at the dorsal-most fin edge in G as indicated by the brackets. **(H)** Single caudal fin bony ray (DIC), somatosensory axons (SS axons), and endothelium of an adult *Tg(p2rx3a>mCherry); Tg(fli1a:EGFP)* fish. Arrowhead denotes a SS axon entering or exiting the bony ray, not associated with endothelial exit points (arrow marks one such example). **(I)** Transverse cross-section of a single bony ray of an adult *Tg(p2rx3a>mCherry); Tg(fli1a:EGFP)* fish. Note the lack of apparent association between axons and endothelium within the intraray space. **(J)** Transverse cross-section of a single bony ray of an adult *Tg(p2rx3a>mCherry)* fish, immunostained with the zns-5 antibody to label osteoblasts. Note the close juxtaposition of axons and intraray osteoblasts. **(K)** Schematic illustrating the anatomy of a transverse view of a single bony ray based on our results. Scale bars, 100 μm (B-G), 50 μm (H), and 25 μm (I,J).

To determine the general organization of peripheral axons within the caudal fin, we stained fins with the monoclonal antibody zn-12 (Metcalfe et al., 1990) and an anti-acetylated tubulin (acTub) antibody, both of which stain multiple types of peripheral axons, and performed high-resolution confocal imaging. In lateral views, we found that axon trajectories followed each of the 18 rays of the caudal fin (Fig. 1B). At ray bifurcations, axon bundles bifurcated to innervate both distal rays (Fig. 1B). Thus, peripheral axons adopt a highly stereotyped pattern mirroring the ray morphology within the adult caudal fin.

### Somatosensory nerves innervate the adult caudal fin rays

Muscles localize only to the base of the caudal fin and a previous analysis based on neuronal dye-filling concluded that motor neurons are largely absent from the caudal fin itself (Schneider and Sulner, 2006). Thus, the peripheral axons we observed likely originated from a different neuronal population. Based on their morphology, we postulated they derived from dorsal root ganglion (DRG) somatosensory neurons.

We took two complementary approaches to establish the identity of axons within the caudal fin. First, we reasoned that mutants with reduced DRG neurogenesis would exhibit decreased fin innervation. To test this prediction, we examined animals homozygous for a predicted null allele of *erbb3b (erbb3b*^*st48/st48*^) (Lyons et al., 2005), in which neural crest migration is disrupted, thereby perturbing DRG neurogenesis (Honjo and Eisen 2008). Consistent with our prediction, staining of *erbb3b* mutant caudal fins for acTub revealed a strong decrease in the amount of bony ray innervation, compared to wild-type fins (Fig. 1C,D). Second, we examined the expression of a reporter for a subset of somatosensory neurons (*Tg(p2rx3a>mCherry)*; (Palanca et al., 2013)) in the caudal fin. We found that the pattern of somatosensory axons recapitulated the ray-associated staining we observed using general axonal markers (Fig. 1E; Supplementary Movie 1). Imaging of this reporter revealed that axons branch within ray segments and exit between dorsal and ventral gaps in the hemirays to innervate the interray space (Fig. 1F, arrowhead). Furthermore, we observed that the rays at the dorsal and ventral margins had a denser concentration of *p2rx3a+* axons than more medial rays (Fig. 1G).

How are somatosensory axons positioned relative to other cell types in the fin? We were particularly interested in the proximity and patterning of axons in relation to endothelium because of their close association in other developmental and tissue contexts (reviewed by Segarra et al., 2019). Through co-imaging of somatosensory axon markers and *Tg(fli1a:EGFP)*, a pan-endothelial reporter (Lawson and Weinstein, 2002), we noted that endothelium and sensory axons both exit through dorsal and ventral gaps in the hemirays, but not generally at the same proximal-distal positions along the rays (Fig. 1H). To establish the lateral axial position of caudal fin axons, we cryosectioned fins and found that somatosensory axons did not obviously associate with endothelial cells located in the center of the intraray space (Fig. 1I). Rather, co-staining with the osteoblast marker zns-5 (Johnson and Weston, 1995), revealed axons adjacent to osteoblasts lining the inner hemiray surface (Fig. 1J; see also Supplementary Movie 1). This localization of somatosensory axons along the concave surface of the inner hemirays is consistent with a previous study that identified neural crest-derived Schwann cells localized to a similar position in the bony rays (Lee et al., 2013). Thus, somatosensory nerves, composed of peripheral axons of DRG neurons and associated Schwann cells, traverse along caudal fin rays in close proximity to osteoblasts that line the inner hemiray surface (Fig. 1K).

### Innervation of the caudal fin rays coincides with endothelial remodeling

To begin to ask how this stereotyped architecture develops, we analyzed the early stages of caudal fin morphogenesis. The caudal fin develops from the larval fin fold, a relatively thin, simple tissue lacking endothelium and bones. The major stages of caudal fin morphogenesis have been described by light microscopy (Bird and Mabee, 2003; Parichy et al., 2009). Briefly, beginning at approximately 4.2 millimeters standard length (mm SL), a mesenchymal condensation forms ventral to the notochord (Fig. 2A, top). Segments of the initial fin rays appear shortly thereafter within the condensation, with pairs of new rays sequentially added on the dorsal and ventral flanks of the initial rays (Fig. 2B-D, top). The first ray joints appear at approximately 6.0 mm SL, with new segments added distally. The fin adopts its characteristic bi-lobed structure beginning around 6.5 mm SL (Fig. 2E, top). Caudal fin endothelial development involves ventral sprouting from posterior axial vessels and formation of a plexus intermediate, which progressively remodels into the mature, repeating pattern of endothelium aligned along and beside the rays (Huang et al., 2009).

**Figure 2.**
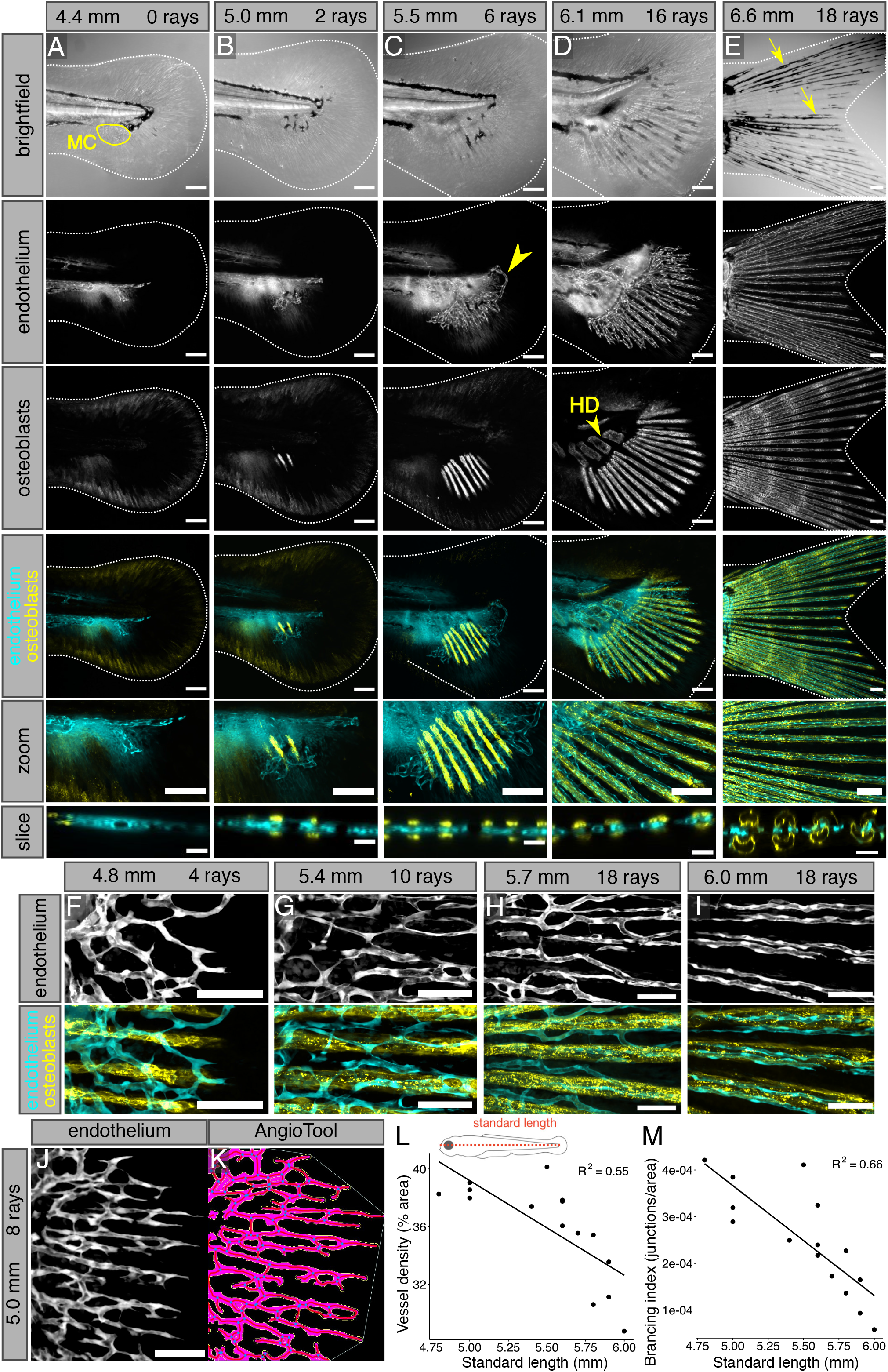
Stages of endothelial outgrowth, plexus formation, and remodeling during caudal fin development. **(A-E)** Confocal projections of caudal fins fixed at the indicated stages expressing transgenic markers for osteoblasts (*Tg(sp7:mCherry-NTR)*) and endothelium (*Tg(fli1a:EGFP)*). Dashed lines denote caudal fin margins. “Zoom” panels reflect cropped, enlarged regions of merged images. “Slice” panels reflect reconstructed orthogonal views of the developing plexus and osteoblasts. The yellow outline in (A) indicates the mesenchymal condensation (MC), visible by brightfield. The yellow arrowhead in C indicates the dorsal endothelial loop. The yellow arrowhead in D indicates the hypural diastema (HD), the future location between hypurals 2 and 3. The yellow arrows in E point to examples of melanophores tracking along bony rays. **(F-I)** Confocal projections of osteoblasts (*Tg(sp7:mCherry-NTR)*) and endothelium (*Tg(fli1a:EGFP)*), capturing endothelial remodeling around the osteoblasts at the indicated stages. Note that the initially disorganized, web-like plexus progressively remodels to its mature linear, ray-aligned morphology. **(J**,**K)** Example of an endothelial (*Tg(fli1a:EGFP)*) image (J), segmented and analyzed in AngioTool (K). Blue dots denote branchpoints. **(L**,**M)** Quantification of density and branching of the caudal fin endothelium relative to standard length. Scale bars, 100 μm (A-E), 20 μm (A-E, slice panels), 50 μm (F-J).

To establish the detailed timing of these stages of endothelial development relative to ray formation, we imaged transgenic animals co-expressing *Tg(fli1a:EGFP)* and *Tg(sp7:mCherry-NTR)*, a reporter for osteoblasts at an intermediate state of differentiation (Knopf et al., 2011; Singh et al., 2012). Prior to the appearance of *sp7+* rays, and concomitant with the formation of the mesenchymal condensation, we observed an initial projection of endothelium into the ventral fin fold (Fig. 2A). This ventral sprout formed at the eventual position of the hypural diastema (HD) (Fig. 2D). The initial two rays formed flanking the dorsal and ventral margins of the endothelial projection, at a stage when the endothelium had begun to form a ventral, fan-shaped plexus (Fig. 2B). At this early stage, we observed blood flowing from the dorsal aorta through the HD and circulating within lumenized vessels in the fin plexus (Supplementary Movie 2), indicating that the fin endothelium had begun to functionally mature. As rays were added sequentially along the dorsal and ventral margins, the plexus continued to expand and eventually connected to the posterior-most axial vessels via a dorsal loop around the 6 ray stage (Fig. 2C, arrowhead; Supplementary Movie 3). Throughout this period, the hemirays flanked the lateral sides of the sheet-like plexus (Fig. 2A-C, slice views).

In contrast to a previous study that proposed that endothelium templates ray growth (Huang et al., 2009), we did not observe the appearance of linear, ray-aligned endothelium at these early stages of fin morphogenesis. Rather, refinement of the vessels to align with the rays occurred between the 14-18 ray stage (Fig. 2F-I). By ∼6.5 mm, when the bi-lobed fin shape emerged, we observed both endothelium and melanophores tracking along the bony rays (Fig. 2E). Quantification of vessel density (percent of vessels per unit area), and branching index (number of vessel junctions per unit area), revealed a negative, linear relationship between each metric and standard length (Fig. 2J-M), highlighting the plasticity of the endothelium during fin organogenesis.

How does the timing of ray innervation relate to the events of bone formation and fin endothelialization that we describe? In one possible model, DRG axons might first pathfind through the developing fin fold tissue, restricting osteogenesis to specific domains, similar to the development of cranial foramina (Akbareian et al., 2015). In another model, osteoblasts might coalesce into hemirays first, through which axons could subsequently pathfind, similar to the radial osteoblast conduits that guide scale innervation (Rasmussen et al., 2018). To distinguish between these possibilities, we visualized axons during the early stages of caudal fin morphogenesis. Using acTub to label all axons in the developing fin, we observed that ray innervation occurred after endothelial sprouting and initial bony ray formation (Fig. 3A-E). Prior to ∼8 rays, the axons in the fin fold tissue were all superficial, arborizing through the epidermis without clear directionality (Fig. 3A,B). Innervation of the rays rapidly increased beginning at ∼10 rays (Fig. 3C). At this stage, we observed bilateral, symmetrical innervation of the developing rays, with axons juxtaposed to *sp7+* osteoblasts found lining the inner hemiray surface (Fig. 3C, slice inset; Supplementary Movie 4). These results suggest that the inner hemiray axon pattern seen in the adult caudal fin is established during the earliest stages of ray innervation. By the 18 ray stage, the fin axons were arranged in a striking candelabra-like pattern (Fig. 3D,E). We measured the development timing of ray innervation against SL and notochord flexion angle, two indicators of maturation, and found that both accurately predicted innervation timing (Fig. 3F,G). In conclusion, the process of caudal fin ray innervation occurs via DRG axons migrating in close association with inner hemiray osteoblasts during a period of active endothelial remodeling.

**Figure 3.**
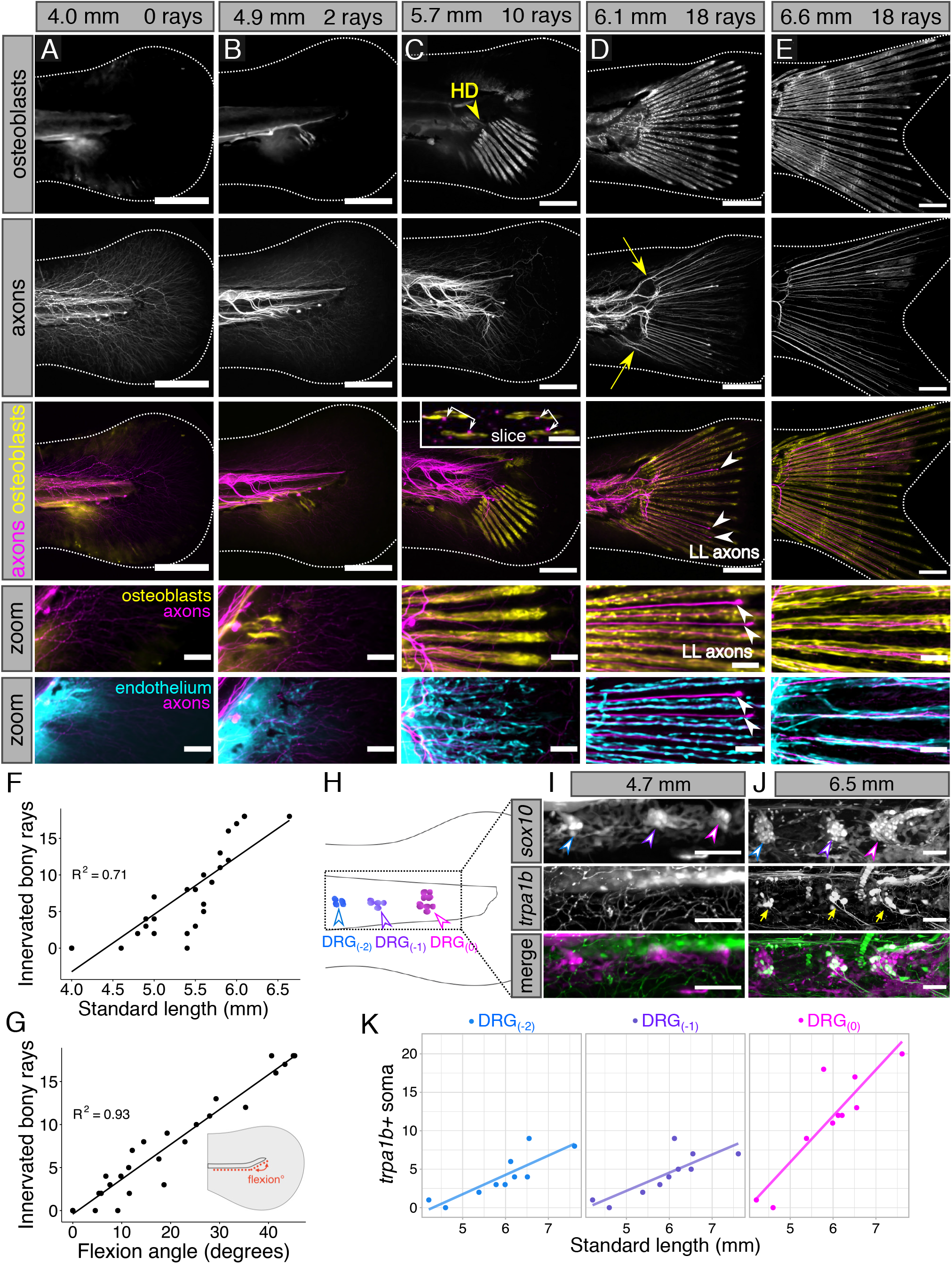
Timing of DRG ray innervation and maturation during caudal fin development. **(A-E)** Confocal projections of caudal fins fixed at the indicated stages expressing transgenic markers for osteoblasts (*Tg(sp7:mCherry-NTR)*) and endothelium (*Tg(fli1a:EGFP)*) and immunostained for axons (acetylated tubulin antibody). “Zoom” images reflect cropped, enlarged regions of merged images. The yellow arrowhead in C denotes the future position of the hypural diastema (HD). New rays are added symmetrically on either side of the HD. Double-headed arrows on the inset in C denote the bilateral, symmetrical pattern of innervation. Note the axons on the inner surface of each hemiray in the slice panel. The yellow arrows in D denote that the dorsal-most and ventral-most ray-associated axons originate along separate routes than the inner ray-associated axons. White arrowheads indicate the position of lateral line (LL) axons growing into the caudal fin in the interray space. **(F**,**G)** Quantification of the number of innervated bony rays versus standard length (F) and notochord flexion angle (G). **(H)** Schematic illustrating the three posterior ganglia of the developing trunk, with DRG_(0)_ (magenta) demarcating the most posterior cluster, DRG_(−1)_ marking the second most posterior cluster (purple), and DRG_(−2)_ marking the third most posterior cluster (blue). **(I**,**J)** Confocal projections of fish expressing neural crest lineage (“*sox10*”: *Tg(Mmu*.*Sox10-Mmu*.*Fos:Cre);Tg(actb2:LOXP-BFP-LOXP-DsRed)*) and sensory channel (*TgBAC(trpa1b:EGFP)*) reporters. Arrowheads as in H. Arrows indicate *trpa1b+* DRG soma. **(K)** Scatterplots with linear regression lines showing quantification of *trpa1b+* soma (arrows in J) in each of the three posterior-most ganglia relative to standard length. Scale bars, 200 μm (A-E), 10 μm (inset in C), and 50 μm (A-E zoom panels, I and J).

### Ray innervation coincides with DRG functional maturation

The initial period of DRG neurogenesis occurs during the first few days of zebrafish development. However, at these early stages, DRG neurons have not been reported to express molecules involved in the detection of somatosensory stimuli, such as the transient receptor potential (TRP) channels Trpa1b or Trpv1 (Esancy et al., 2018; Gau et al., 2013; Pan et al., 2012; Prober et al., 2008). How might the period of fin innervation we identified relate to the functional maturation of DRG neurons? To address this question, we examined the expression of a reporter for *trpa1b* (*TgBAC(trpa1b:EGFP)*; (Pan et al., 2012)), which encodes a TRP channel that detects noxious chemicals (Prober et al., 2008), in the three most posterior dorsal root ganglia in the trunk (Fig. 3H). By quantifying *trpa1b+* neuronal soma within these ganglia, we found that the onset of *trpa1b* expression tightly corresponded to the period of active fin innervation (Fig. 3I-K). Thus, DRG nerves not only pathfind through the bony rays at early stages of caudal fin organogenesis, but they also begin to express functional sensory molecules.

### *Caudal fin endothelialization* requires *VEGFR signaling*

To test for an interdependence between tissue types during caudal fin development, we first sought to identify molecular signals required for fin endothelialization. Due to its function regulating angiogenesis during caudal fin regeneration (Bayliss et al., 2006), we considered vascular endothelial growth factor receptor (VEGFR) a strong candidate. To block VEGFR signaling, we used PTK787, a highly specific pharmacological inhibitor of VEGFR (Bayliss et al., 2006). Compared to DMSO-treated controls, we found that application of 0.5 μM PTK787 beginning at 4.2-4.4 mm SL completely blocked endothelial growth in the developing fin (Fig. 4A and 4B-G, top). Despite PTK787 systemically inhibiting VEGFR, fish without caudal fin endothelium progressed through similar developmental metrics of maturation, including notochord flexion, standard length, and number of bony rays, compared to controls (Fig. 4H,I). However, we did note that PTK787-treated fins plateaued with an upper size limit of ∼1 mm^2^ (Fig. 4J). Thus, VEGFR signaling is necessary for caudal fin endothelialization and the rapid expansion of fin size that occurs during organogenesis.

**Figure 4.**
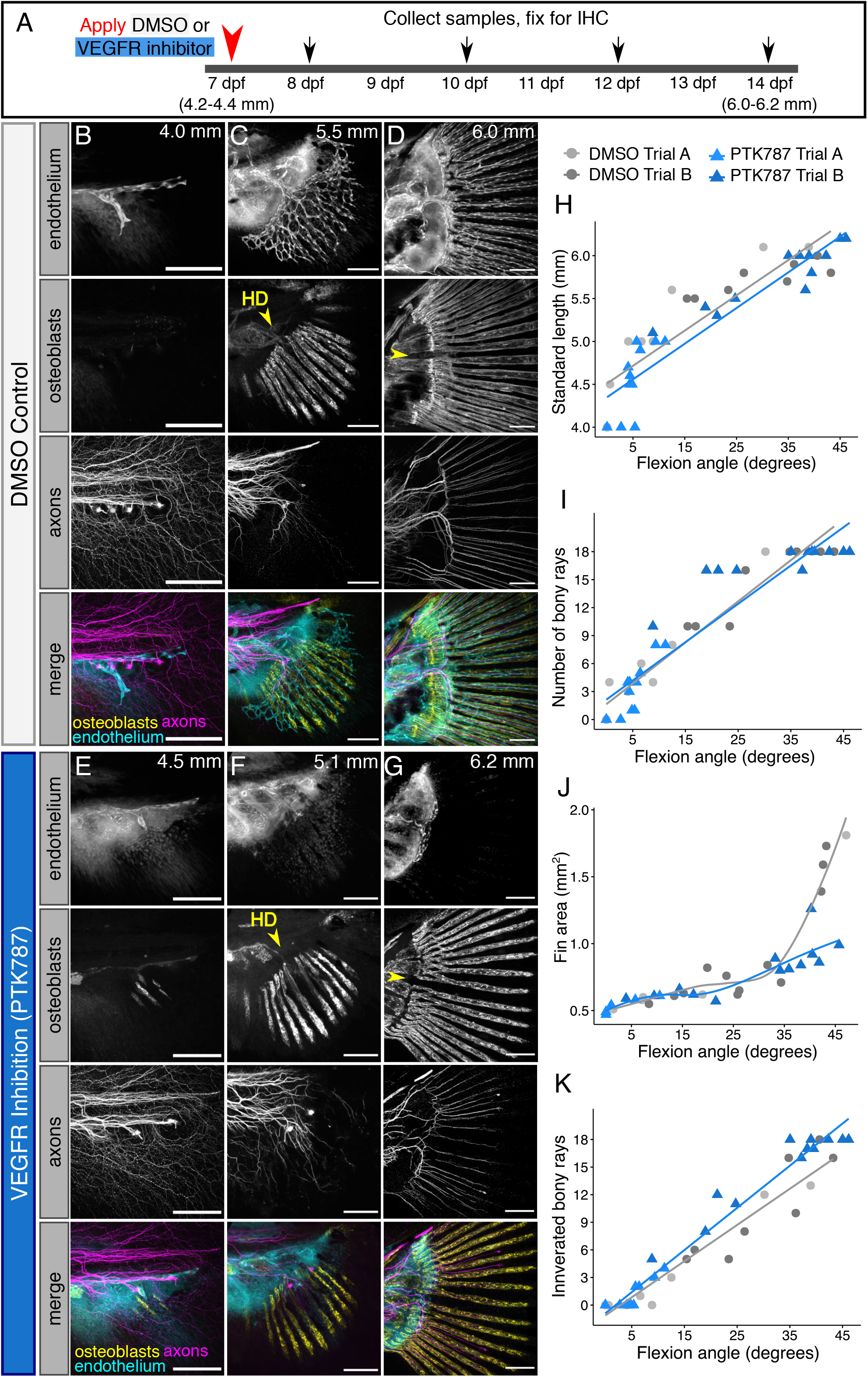
Caudal fin bony ray development and innervation progress in the absence of endothelium. **(A)** Diagram illustrating experimental scheme of VEGFR inhibition in developing zebrafish. **(B-G)** Confocal projections of fins expressing reporters for osteoblasts (*Tg(sp7:mCherry-NTR)*) and endothelium (*Tg(fli1a:EGFP)*) and immunostained for axons (acetylated tubulin antibody). Treatments and stages as indicated. Yellow arrowhead denotes the future position of the hypural diastema (HD). Note that despite the severe reduction in endothelial growth into the fin following PTK787 treatment, early osteoblast and axon patterning appear unperturbed. **(H-K)** Quantification of developmental metrics and ray innervation between treatment groups. Lines represent linear regression for (H,I,K) and local polynomial smoothing for (J). SL/flexion angle slope between the two independent trials of DMSO control fish and PTK treated fish was not significantly different (*P* = 0.158; ANCOVA). Flexion/number of bony rays slope between the two independent trials of DMSO control fish and PTK treated fish was not significantly different (*P* = 0.913; ANCOVA). Number of innervated rays/flexion angle slope between the two independent trials of DMSO control fish and PTK treated fish was significantly different above a flexion angle of 13.06 degrees (*P* <0.05; Johnson-Neyman technique). Scale bars, 100 μm (B-G).

### Endothelium is not required for initial fin ray and nerve morphogenesis

We next sought to test the requirement of each respective tissue—endothelium, osteoblasts, and DRG neurons—on each other’s development during fin maturation. Previous work proposed that endothelium guides osteoblast patterning during caudal fin morphogenesis (Huang et al., 2009), however, this requirement was not formally tested. To directly assess the requirement of endothelium in tissue patterning during fin morphogenesis, we treated animals with PTK787 to inhibit endothelialization of the fin and examined osteoblast and axon organization (Fig. 4A). Remarkably, when we examined *sp7+* osteoblasts in PTK787-treated fins, we observed that the osteoblasts continued their symmetrical bony ray outgrowth, similar to controls (Fig. 4B-G). In the absence of an endothelial plexus, a full suite of 18 bony rays formed by 6 mm SL and appeared to initiate the process of segmentation, similar to controls (Fig. 4D,G and data not shown). Vessels influence the development and homeostasis of peripheral nerves in many contexts (reviewed by Segarra et al., 2019). To determine whether ray innervation requires endothelium, we stained PTK787-and DMSO-treated fish for acTub. We found that the pattern and timing of DRG axons innervating the bony rays were comparable between PTK787-treated and controls (Fig. 4B-G,K). Thus, we conclude that endothelium is not required for osteoblast or axon patterning during the early stages of caudal fin organogenesis.

### Ray-associated osteoblasts and endothelium pattern independently of DRG neurons

Peripheral nerves provide cues that pattern vascular development in mouse limb skin (Li et al., 2013; Mukouyama et al., 2002; Mukouyama et al., 2005). While nerves are required for pectoral fin regeneration (Geraudie and Singer, 1985; Simões et al., 2014), their role in regulation of fin organogenesis has yet to be investigated. To assess the potential contributions of DRG nerves to fin development, we examined the integrity of caudal fin bony rays and endothelium in mutants whose DRG neurons do not properly develop (*erbb3b*^*st48/st48*^ or *adgra2*^*s984/s984*^ mutants; (Honjo et al., 2008; Vanhollebeke et al., 2015)). As predicted based on our observations in adults, *adgra2* and *erbb3b* mutant fins completely lacked ray-associated innervation (Fig. 5A-C) and had significantly reduced fin axon density compared to sibling controls (Fig. 5D). We imaged bony ray calcification with Alizarin red staining and did not visually identify any aberrations in early ray patterning relative to sibling controls (Fig. 5A-C). We next evaluated the endothelial patterning in the caudal fins of these mutant fish and did not observe significant differences between siblings and mutants for either genotype, except for the number of endpoints between *erbb3b* mutants and siblings (Fig. 5E-F). Our observations indicate that DRG neurons are largely dispensable for endothelial and osteoblast patterning during the early stages of fin organogenesis.

**Figure 5.**
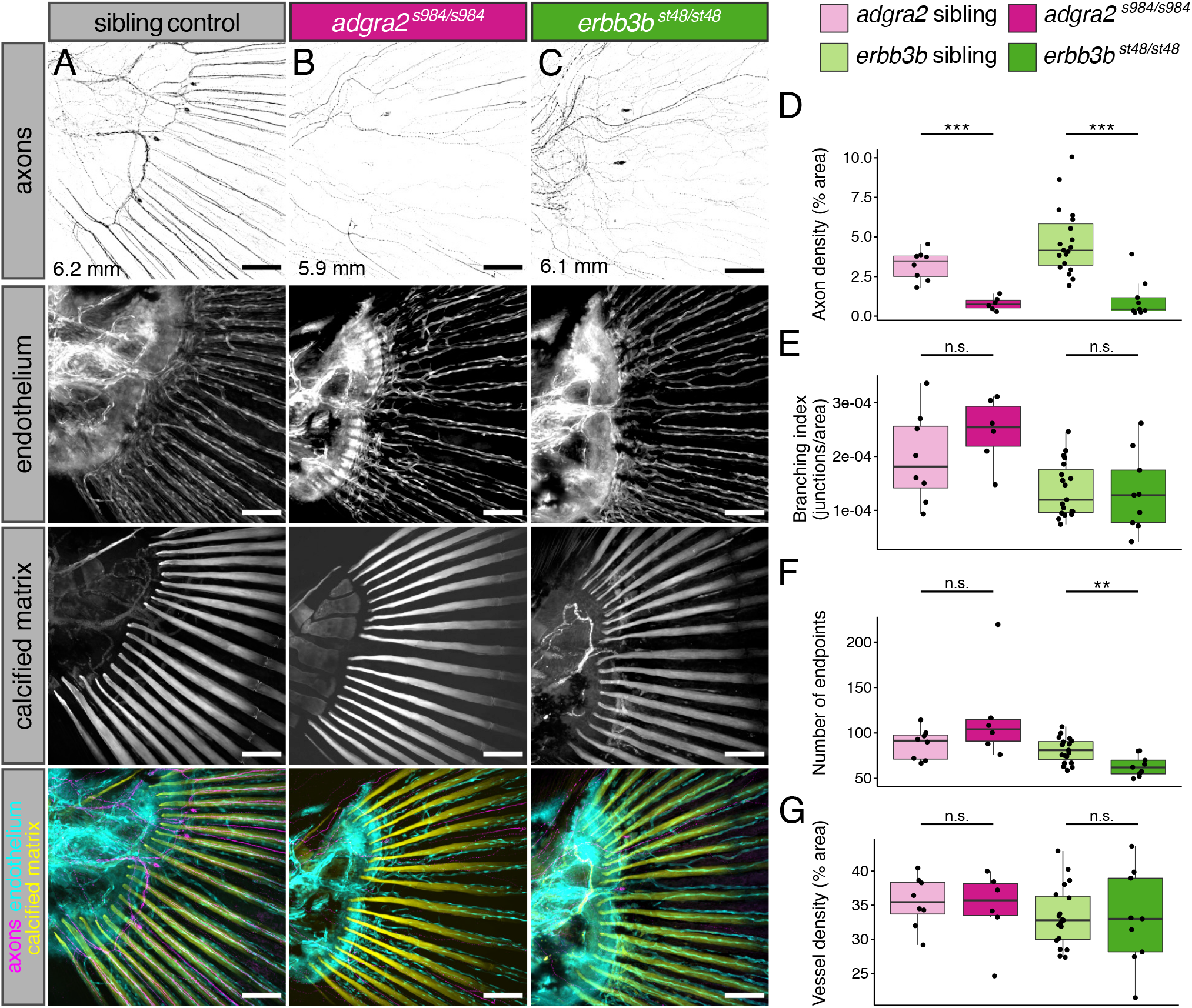
Innervation of the caudal fin bony rays by DRG axons is not required for caudal fin ray and endothelial patterning. **(A-C)** Representative confocal projections of caudal fins from the indicated genotypes expressing the endothelial marker *Tg(fli1a:EGFP)* fixed at the indicated stages and stained for axons (acetylated tubulin antibody) and calcified matrix (Alizarin red). Note the lack of ray-associated axons in the *adgra2* and *erbb3b* mutants. **(D-G)** Boxplots of axon density, branching index, number of endpoints, and vessel density for fins of the indicated genotypes. Wilcoxon rank sum test between *adgra2* siblings and *adgra2* mutants (axon density, *P* = 0.000666; branching index, *P* = 0.345; number of endpoints, *P* = 0.1551; vessel density, *P* = 0.8518). *adgra2* sibling fish, *n* = 8 (mean SL = 6.01 mm ± 0.11 s.d.). *adgra2* mutant fish, *n* = 6 (mean SL = 6.0 mm ± 0.48 s.d.). Wilcoxon rank sum test between *erbb3b* siblings and *erbb3b* mutants (axon density, *P* = 2.809e-05; branching index, *P* = 0.6993; number of endpoints, *P* = 0.004654; vessel density, *P* = 0.9615) *erbb3b* sibling fish, *n* = 19 (mean SL = 6.34 mm ± 0.29 s.d.). *erbb3b* mutant fish, *n* = 9 (mean SL = 6.89 mm ± 0.45 s.d.). * indicates *P*<0.05, ** indicates *P*<0.01, *** indicates *P*<0.001. Scale bars, 100 μm (A-C).

### sp7+ osteoblasts are required for endothelial remodeling and axon outgrowth

Informed by our previous findings that demonstrated a role for osteoblasts in guiding nerves and endothelium in developing scales (Rasmussen et al., 2018), we next asked if the bony rays themselves established the pattern of caudal fin endothelium and innervation. To determine whether endothelial or axon development require osteoblasts, we took advantage of a previously described transgenic strategy to inducibly ablate *sp7+* osteoblasts during the early stages of caudal fin morphogenesis. The transgenic line *Tg(sp7:mCherry-NTR)* expresses the nitroreductase enzyme (NTR) fused to mCherry, which allows for conditional ablation of *sp7+* osteoblasts upon addition of metronidazole (MTZ) (Singh et al., 2012).

To ablate *sp7+* osteoblasts in the developing caudal fin, we applied 9 mM MTZ to double transgenic *Tg(sp7:mCherry-NTR);Tg(fli1a:EGFP)* fish beginning at 4.2-4.4 mm SL, or ∼7 dpf (Fig. 6A). After an initial 24 h exposure, we then maintained fish in 4.5 mM MTZ for the remainder of the experimental trial, to avoid MTZ-mediated toxicity. As controls, we exposed single transgenic *Tg(fli1a:EGFP)* siblings to the same MTZ treatment. To confirm that we had ablated osteoblasts, we stained treated fins with zns-5 and Alizarin red, which stains calcified matrix. We observed complete elimination of zns-5+ osteoblasts and calcified bone compared to controls (Fig. 6B,C). To account for any effects of our ablation strategy on animal or organ growth, we compared developmental metrics between treatment groups. We found that while the overall fin area of *sp7+* ablated fish was significantly decreased compared to control treated fish, metrics of maturation (flexion angle, SL) were not significantly different between the two groups (Fig. 6D,E). Thus, our ablation regimen efficiently ablated *sp7+* osteoblasts, while having minimal effects on overall larval development.

**Figure 6.**
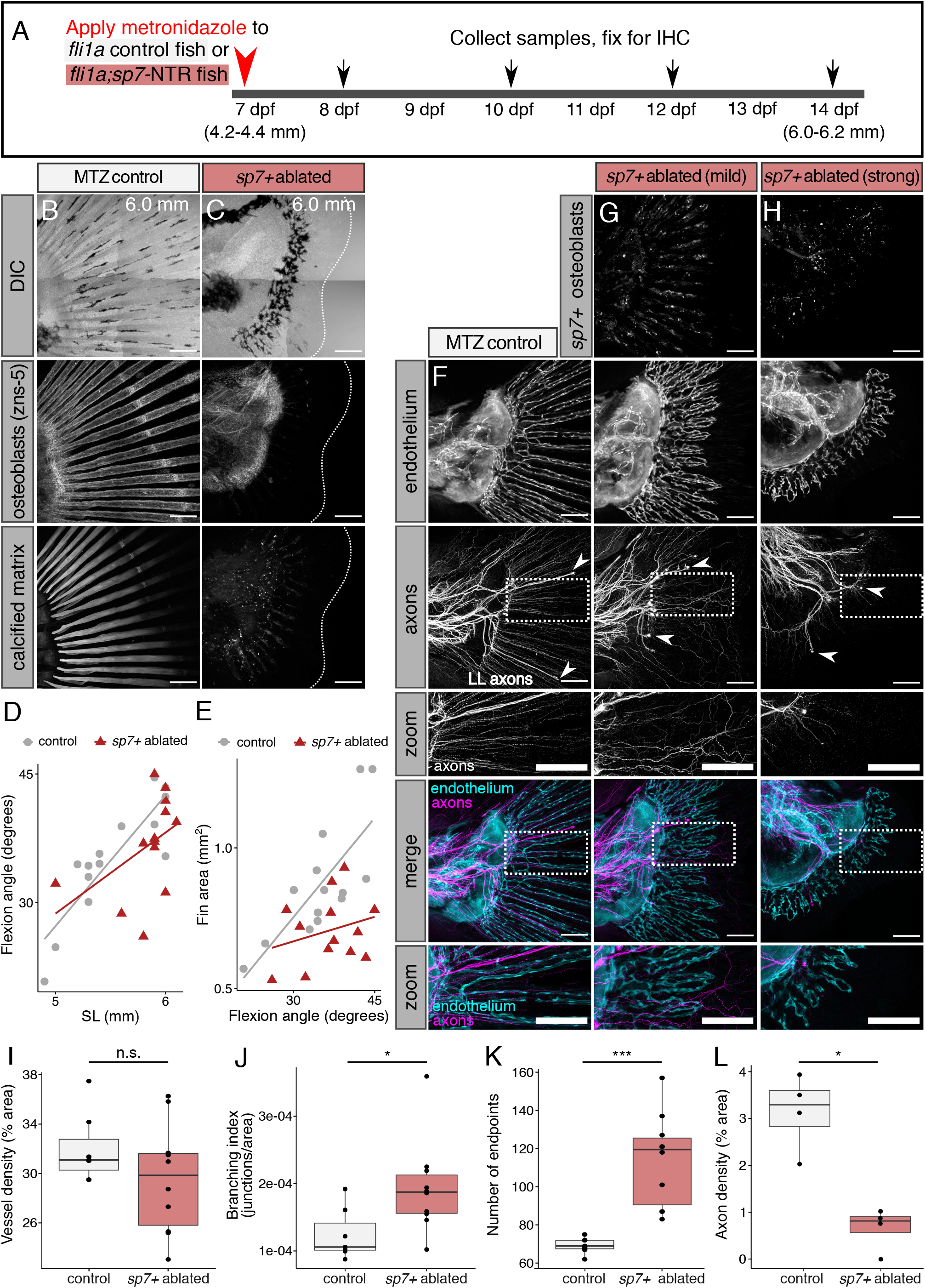
Osteoblasts are required for endothelial remodeling and axon patterning in the developing caudal fin. **(A)** Diagram illustrating experimental scheme of *sp7+* osteoblast ablation in developing zebrafish. **(B**,**C)** Representative confocal projections of MTZ-treated control (*Tg(fli1a:EGFP)*) or osteoblast-ablated (*Tg(fli1a:EGFP);Tg(sp7:mCherry-NTR)*) caudal fins fixed at the indicated stages and stained for a marker of osteoblasts (zns-5 antibody) and calcified matrix (Alizarin red). Dashed lines denote caudal fin margins. Note the lack of zns-5+ osteoblasts and calcified matrix within the osteoblast ablated fin. **(D**,**E)** Developmental metrics in control and *sp7+* ablated fins. Control *fli1a+* fish, *n* = 14. *sp7+* ablated fish, *n* = 13. SL/flexion slope between genotypes was not significantly different (*P* = 0.154; ANCOVA). Flexion/fin area between genotypes was significantly different (*P* = 0.00102; ANCOVA). Representative results from a single independent trial. **(F-H)** Representative confocal projections of MTZ-treated control (*Tg(fli1a:EGFP)*) or osteoblast-ablated (*Tg(fli1a:EGFP);Tg(sp7:mCherry-NTR)*) caudal fins fixed at the indicated stages and stained for axons (acetylated tubulin antibody). Yellow arrowheads indicate lateral line (LL) axons. Note the disorganized appearance of axons and endothelium in the absence of *sp7+* osteoblasts. **(I-K)** Boxplots of vessel density, branching index, and number of endpoints between treatment groups. Wilcoxon rank sum test results: Vessel density, *P* = 0.31; branching index, *P* = 0.033; number of endpoints, *P* = 0.00074. Control *fli1a+* fish, *n* = 7 (mean SL = 5.96 mm ± 0.05 s.d.). *sp7+* ablated fish, *n* = 10 (mean SL = 5.87 mm ± 0.15 s.d.). Representative results from a single independent trial. **(L)** Boxplot of axon density between control and osteoblast-ablated groups. Wilcoxon rank sum test between groups, *P* = 0.02857. Control *fli1a+* fish, *n* = 4 (mean SL = 5.9 mm ± 0.00 s.d.). *sp7+* ablated fish, *n* = 4 (mean SL = 5.93 mm ± 0.09 s.d.). * indicates *P*<0.05, *** indicates *P*<0.001. Scale bars, 100 μm (B,C,F-H).

We evaluated endothelial and nerve patterning in MTZ-treated fins and found that *fli1a+* vessels grew into developing fins in the absence of *sp7+* osteoblasts (Fig. 6F-H). Quantitative analysis of vessel patterning revealed that vessel density was not significantly different in the *sp7+* ablated fins compared to controls (Fig. 6I). However, treated fins did exhibit a significant increase in the number of vessel endpoints (fragmentation) and vessel branching compared to controls (Fig. 6J,K). Additionally, we observed that in the absence of *sp7+* osteoblasts, DRG axon pathfinding was stunted and disorganized and innervation density was significantly decreased (Fig. 6G,H,L). Together, these results suggest that while *sp7+* osteoblasts are not required for the initial events of caudal fin organogenesis, including ventral endothelial sprouting and plexus formation, they are required for the later stages of endothelial remodeling, axon innervation, and organ growth.

## Discussion

Congruent neurovascular-bone patterning has been documented in diverse tissues for over 150 years, yet how this congruence is generally achieved remains an unanswered question in vertebrate development. In this study, we pioneer the use of the zebrafish caudal fin—an optically clear and relatively two-dimensional organ—for developmental analysis of neurovascular-bone interactions. Despite being extensively used as a model for organ regeneration (reviewed by Pfefferli and Jaźwińska, 2015; Poss et al., 2003), the identity of peripheral nerves within the caudal fin, their relationships to other tissue types, and the mechanisms of their innervation have not been analyzed in-depth. Here, we directly address these knowledge gaps and establish a cellular hierarchy during fin development. Based on our results from systematically blocking the development of bones, vessels, and sensory neurons, we propose that osteoblasts function as central organizers that pattern the two other cell types during fin organogenesis (Fig. 7).

**Figure 7.**
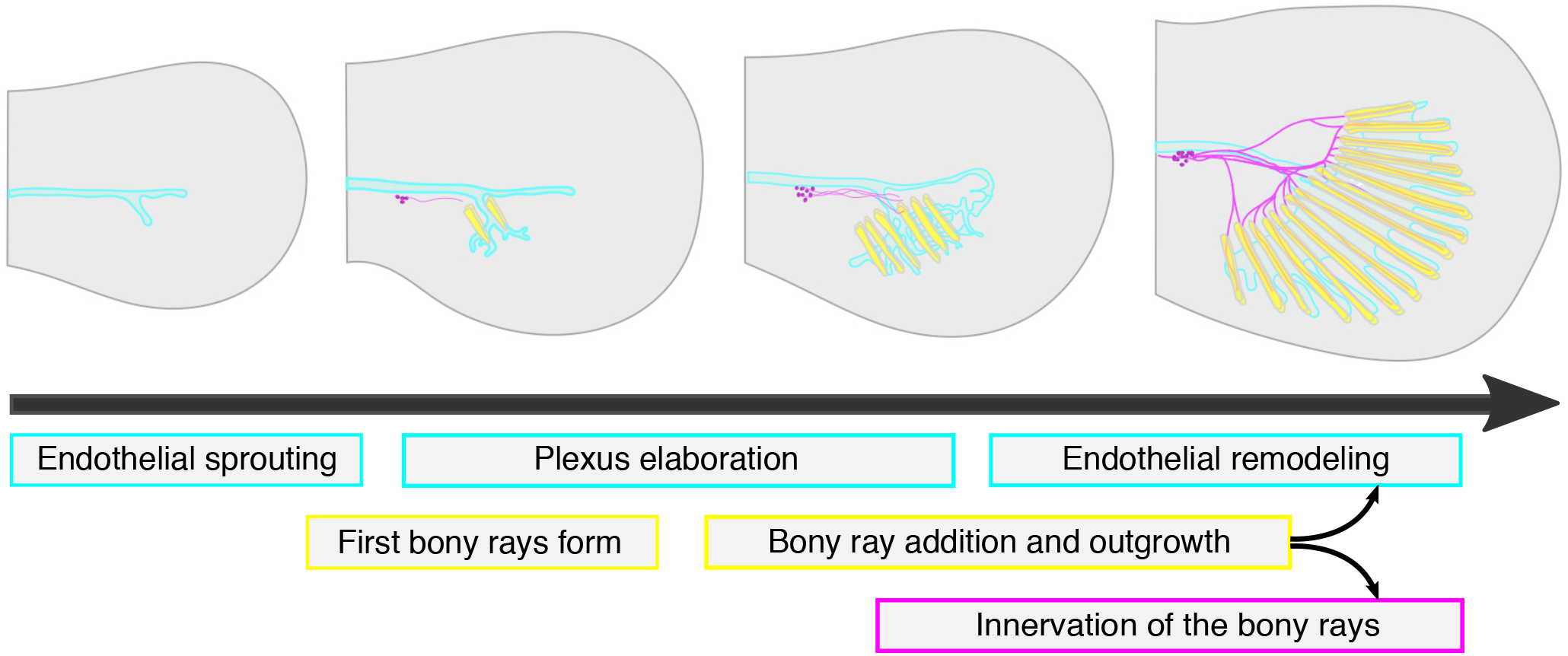
Summary of caudal fin ray, endothelium, and axon development. Schematic illustrating the sequential cellular events between caudal fin rays, endothelium, and axons during caudal fin development described in this study. Arrows indicate dependence of endothelial remodeling and fin innervation on the presence of *sp7+* osteoblasts along the rays.

The tight association between nerves and ray-associated osteoblasts that we report here in the zebrafish caudal fin extends previous findings in several teleosts. Histological studies from the 1980s reported the “collagen fibrils” of nerve fibers within the caudal fin bony rays of tilapia (Becerra et al., 1983; Montes et al., 1982). Recent work in medaka, another teleost, identified nerves running through the bony rays and co-localizing with myelinating cells (Dodo et al., 2020). Further, axon-associated neural crest-derived Schwann cells localize to the inner hemiray surface in zebrafish (Lee et al., 2013). Using somatosensory-specific reporters and mutant analysis, we show that the axon bundles at the inner, concave osteoblast-lined surface are the peripheral axons of DRG neurons. We hypothesize that these sensory axons utilize the bony ray tracts before innervating surrounding fin tissue as protective “armor” to avoid injury and debilitating damage. We further find that different fin regions have variable densities of somatosensory innervation, akin to anatomical differences observed in humans (Johansson and Vallbo, 1979). Together, our work and others demonstrate that somatosensory axons traverse the relatively long distance of the adult caudal fin by running through the dermal bony rays, tightly juxtaposed to intraray osteoblasts (Fig. 1K), and not the intraray arteries, as previously suggested (Tu and Johnson, 2011).

Our developmental analysis showed that caudal fin morphogenesis—specifically, the development of its bone, nerves, and endothelium—is a rapid process of sequential outgrowth, proliferation, interaction, and remodeling. An initial endothelial projection invades the ventral side of the fin fold, followed rapidly by the formation of the first paired bony rays (Fig. 7). As bony rays are added sequentially and symmetrically around the eventual hypural diastema, the endothelial plexus continues its outgrowth as a dense, web-like network (Fig. 7). Beginning around the 8-10 bony ray stage, axons pathfind through the inner surface of the paired hemirays (Fig. 7). The fin then undergoes rapid proliferation and extension of all three tissue types; osteoblasts deposit a full suite of 18 bony rays, endothelium remodels around the bony ray tracts, and axon bundles race down the inner hemiray surfaces towards the distal end of the fin. The ability to visually record these processes highlights the strength of the zebrafish caudal fin as a powerful *in vivo* model to explore the cellular basis of the neurovascular-bone relationship.

Previous studies suggest that the axons we observed arborizing within the early fin fold (e.g, Fig. 3A) are likely Rohon-Beard (RB) neurons (Rieger and Sagasti, 2011), a transient population of somatosensory neurons in anamniotes. As the fish mature from larval to juvenile stages, there is a transition in trunk and fin innervation from RB to DRG neurons (Rasmussen et al., 2018; Williams and Ribera, 2020). In contrast to the freely branching RB axons, we observed that DRG axons innervating the fin adopted a stereotyped, candelabra-like morphology beginning around the 8-10 ray stage. Although an initial population of DRG neurons develops by 3 dpf and expresses molecules involved in somatosensory neuron specification such as *neurogenin 1* (Andermann et al., 2002; Cornell and Eisen, 2002), at these early stages DRG neurons are not reported to express the nociceptive markers *trpa1b* or *trpv1* (Esancy et al., 2018; Gau et al., 2013; Pan et al., 2012; Prober et al., 2008). We describe the concurrent timing of bony ray innervation and expression of *trpa1b* in a subset of DRG neurons. Whether these *trpa1b+* neurons reflect maturation of pre-existing neurons or addition of newly born neurons, which has been previously documented during these stages (McGraw et al., 2012), remains unknown. Our analysis of mutants lacking DRG neurons demonstrated that neither endothelial nor ray patterning require somatosensory nerves. These results contrast with chick and mouse limb skin, where innervation induces local vascular remodeling (Li et al., 2013; Martin and Lewis, 1989; Mukouyama et al., 2002; Mukouyama et al., 2005).

Huang et al. (2009) proposed that the caudal fin endothelium guides osteoblasts, thereby templating the subsequent bony rays they form. Moreover, a three-dimensional superstructural analysis demonstrated continuous intercellular communication pathways between vasculature and developing caudal fin rays (Akiva et al., 2019). While we observed active blood flow in the fin at the stages of ray development and ray innervation (Supplementary Movies 2 and 3), blocking endothelialization with the VEGFR inhibitor PTK787 did not inhibit either of these processes. Thus, although active blood flow through the endothelial plexus is an early feature of caudal fin development, endothelial tissue is not required for the initial morphogenesis of caudal fin bony rays or their innervation by somatosensory axons. Nevertheless, we did find that PTK787-treated fins had a smaller overall size, suggesting that, similar to observations during fin regeneration (Bayliss et al., 2006), the avascular organ is limited in developmental growth potential. Further experiments are required to determine whether the limitation in organ growth in the absence of endothelium reflects hypoxic conditions or a necessity for endothelial signaling.

In our study, we examined the requirement of *sp7+* osteoblasts in patterning caudal fin endothelium and axons. It is important to note that our ablation strategy did not target *runx2+* pre-osteoblasts or collagen-rich actinotrichia that localize to the leading edge of bony rays (Brown et al., 2009; Durán et al., 2011). Thus, with ray tips presumably intact, we perturbed only the intermediate osteoblast population (Fig. 6) and observed disrupted endothelium and axon patterning, but not complete disorganization. This finding suggests that *sp7+* osteoblasts are critical to the process of endothelial plexus remodeling, but not initial outgrowth. Our results are consistent with recent work in the pectoral fin that observed a tight temporal relationship between endothelial remodeling and bony ray formation and suggested that bone cells likely produce proangiogenic cues (Paulissen et al., 2021). In addition, since we observed wandering and stunted axon patterning following osteoblast ablation, our results suggest that *sp7+* osteoblasts may also be required to produce essential neurotrophic signals to attract sensory axon growth cones. One limitation to our approach is that MTZ-based ablation may have cytotoxic or systemic effects. However, our observation that overall vessel density was not significantly different following osteoblast ablation (Fig. 6I) would argue against non-specific effects. Future improvement in ablation techniques and specificity, as well as additional molecular analyses, will build on our findings.

Nerve-dependent regeneration has been observed in response to appendage amputation in diverse animal phyla, from axolotl limbs (Kragl et al., 2009), to spiny mouse skin (Seifert et al., 2012), to zebrafish pectoral fins (Simões et al., 2014). While the zebrafish caudal fin is a well-established model of appendage regeneration (reviewed by Pfefferli and Jaźwińska, 2015; Poss et al., 2003), the role that sensory nerves play in its repatterning is not yet clear. Intriguingly, inhibition of cholinergic signaling in the caudal fin inhibits regeneration (Recidoro et al., 2014), suggesting that neuronal-derived cues are likely required. Dissecting the cellular and molecular mechanisms of bone-associated innervation during early caudal fin development may shed light on the processes that enable organ re-innervation after severe injury.

## Materials and Methods

### Zebrafish

Zebrafish (*Danio rerio)* were maintained at 26-27°C on a standard 14 h light/10 h dark cycle. Fish were staged by standard length (SL) (Parichy et al., 2009). Larval fish were raised in 0.8 L tanks at a maximum density of 50 fish per tank. Larvae were fed rotifers once or twice a day at a rate of 25,000-50,000 per tank until 10 days post-fertilization (dpf), after which they were fed artemia in addition to rotifers. SL of adult fish under anesthesia was measured with a standard ruler before performing caudal fin biopsies. SL of larval fish was measured using the IC Measure software (The Imaging Source) on images captured on a Stemi 508 stereoscope (Zeiss) equipped with a DFK 33UX264 digital camera (The Imaging Source). Animals >18 mm SL were used for adult portions of the study (Fig. 1). For larval studies, notochord flexion angle was also recorded as a marker of developmental progression (Parichy et al., 2009). All animal husbandry and experimental procedures were approved by the University of Washington Office of Animal Welfare (protocol #4439-01).

The following strains were used: AB (Wild-Type), *Tg(Ola*.*Sp7:mCherry-Eco*.*NfsB)*^*pd46*^ (referred to as *Tg(sp7:mCherry-NTR)*; (Singh et al., 2012)), *Tg(sp7:EGFP)*^*b1212*^ (DeLaurier et al., 2010), *Tg(fli1a:EGFP)*^*y1*^ (Lawson and Weinstein, 2002), *Tg(Tru*.*P2rx3a:LEXA-VP16,4xLEXOP-mCherry)*^*la207*^ (referred to as *Tg(p2rx3a>mCherry)*; (Palanca et al., 2013)), *TgBAC(trpa1b:EGFP)*^*a129*^ (Pan et al., 2012), *Tg(Mmu*.*Sox10-Mmu*.*Fos:Cre)*^*zf384*^ (Kague et al., 2012), *Tg(actb2:LOXP-BFP-LOXP-DsRed)*^*sd27*^ (Kobayashi et al., 2014), *Tg(−17*.*0neurog1:EGFP)*^*w61*^ (McGraw et al., 2008), *erbb3b*^*st48*^ (Lyons et al., 2005), and *adgra2*^*s984*^ (Vanhollebeke et al., 2015).

### Whole mount fixation

Samples were fixed in 4% paraformaldehyde in phosphate-buffered saline (PBS) overnight at 4°C. After fixation, samples were washed and permeabilized following an established protocol (Huang et al., 2009). Briefly, samples were washed 2×5 min in PBS, then 1×5 min in H2O. They were permeabilized with acetone for 7 min at −20°C, followed by a 1×5 min wash in H2O and a 1×5 min wash in PBS.

### Whole mount staining

Samples were blocked in 5% goat serum in PBST (PBS + 0.3% Triton) for at least 1 h at room temperature prior to antibody staining (König et al., 2019), then incubated in primary antibody diluted in PBST overnight at 4°C. The following day, following 4×15 min washes in PBST, samples were incubated with the appropriate secondary antibody diluted in PBST overnight at 4°C. Samples were washed 4×15 min in PBST, before incubation in DAPI (5 ng/μl). Samples were mounted on a glass slide in ProLong Gold antifade (ThermoFisher) and a coverslip was applied. The following antibodies were used: rabbit monoclonal anti-acetylated alpha tubulin (Cell Signalling Technology; RRID:AB_10544694; 1:800), mouse monoclonal zns-5 (Zebrafish International Resource Center; RRID:AB_10013796; 1:200), mouse monoclonal zn-12 (Zebrafish International Resource Center; RRID:AB_10013761; 1:200), Goat anti-Rabbit IgG (H+L) Cross-Adsorbed Secondary Antibody, Alexa Fluor 647 (ThermoFisher; RRID:AB_2633282; 1:500), and Goat anti-Mouse IgG (H+L) Cross-Adsorbed Secondary Antibody, Alexa Fluor 488 (ThermoFisher; RRID:AB_2534069; 1:500).

### Transverse cross sections

Caudal fin biopsies were fixed in 4% formaldehyde in PBS at room temperature for 1 h. Fins were then subjected to a sucrose sink and embedding steps, following an established protocol (Petrie et al., 2014). Briefly, tissue was washed in 5% sucrose in PBS for 30 min at room temperature. Then, fins were washed 2×1 h in 5% sucrose in PBS. The solution was changed to 30% sucrose in PBS and samples were placed at 4°C on a nutator overnight. Finally, sucrose solution was adjusted to a 1:1 ratio of 30% sucrose:100% O.C.T. compound (Tissue-Tek, VWR #25608-930) and placed at 4°C on a nutator overnight. Tissue was then directly embedded in 100% OCT in embedding wells and stored at −80°C.

Sectioning was performed on a Leica CM1850 cryostat. Fins were sectioned in the transverse orientation at 18 µm thickness. Sectioned samples were adhered to positively charged slides and stored at -80°C until downstream processing. Retrieved slides were placed on a heat block at 37°C for 30 min to evaporate moisture. A hydrophobic pen (Cole-Parmer Hydrophobic Barrier PAP Pen # UX-75955-53) was used to carefully trace an outline of samples and left to dry for 10-15 min. All staining, blocking, and washing was performed as described above, but in a moisture chamber. After the final wash, a small drop of antifade and a coverslip were applied.

### Live imaging of larval fish

Fish were anesthetized in 100 μg/ml tricaine (MS-222) in system water until no longer motile. Animals were mounted in a glass bottom dish in 0.6% low melt agarose dissolved in system water. Insect pins were used to gently manipulate the fish into place. Once the agarose had set, tricaine solution was added to the dish for imaging. After imaging was complete, fish were removed from the agarose and monitored in fresh system water until swimming again.

### Live imaging of blood flow

To visualize blood flow in the developing caudal fin, larval fish were anesthetized and mounted as described above. Differential interference contrast (DIC) images were captured on a A1R-MP+ confocal microscope (Nikon), using continuous image acquisition at a rate of 15 frames per sec.

### Vascular growth inhibition

The VEGFR inhibitor PTK787 (Tocris Bioscience; Cat# 5680) was used to inhibit angiogenesis as previously described (Chan et al., 2002). PTK787 was dissolved in DMSO to a stock concentration of 50 mM. Treatments were performed on cohorts of 25 *Tg(fli1a:EGFP); Tg(sp7:mCherry-NTR)* fish per condition. Fish were treated with either 0.5 μM PTK787 or 0.00001% DMSO in system water. Fish were housed in 400 mL of static system water for the duration of the study. Regular feeding of rotifers and artemia was performed, and 80% of the water was exchanged every 2 d with fresh DMSO or PTK787 solution. Treatment began before the visible appearance of *sp7+* osteoblasts in the caudal fin (approximately 4.2-4.4 mm SL, or ∼7 dpf). Fish were maintained in the drug for a maximum of 7 d before being sacrificed for analysis. Standard length, flexion angle, and overall fin area were measured to evaluate potential developmental delay between treatment groups.

### Osteoblast ablation

Ablation of osteoblasts was performed with a chemically inducible, genetic approach. *Tg(sp7:mCherry-NTR)* fish (Singh et al., 2012) express nitroreductase (NTR) under the intermediate osteoblast promoter, *sp7* (previously known as *osterix/osx)*. NTR renders the prodrug metronidazole (MTZ) cytotoxic, killing all *sp7+* osteoblasts after drug application. For this study, an initial dose of 9 mM MTZ (Sigma Aldrich; Cat# M1547) was applied at approximately 4.2-4.4 mm SL, or ∼7 dpf. After 24 h, fish were maintained at a 50% dose (4.5 mM) for the remainder of the experimental trial. Water and MTZ were exchanged every 48 h. This drug regimen did not result in significant morbidity. Sibling fish lacking *Tg(sp7:mCherry-NTR)* were subjected to the same drug treatment. Standard length, flexion angle, and overall fin area were measured using the IC Measure software and recorded to evaluate potential developmental delay between experimental groups. Alizarin red staining (marking calcified matrix; ACROS Organics; Cat# 400480250) and zns-5 antibody staining (alternative marker of osteoblasts) were used to confirm complete ablation of osteoblasts.

### DRG mutant identification

*erbb3b*^*st48/+*^ and *adgra2*^*s984/+*^ adults were identified as described (Lyons et al., 2005; Vanhollebeke et al., 2015). Heterozygous adults were incrossed and homozygous mutant progeny were identified based on a reduction in larval dorsal root ganglion neurons (Honjo et al., 2008; Vanhollebeke et al., 2015) using *Tg(−17*.*0neurog1:EGFP)* as a marker (McGraw et al., 2008).

### Confocal imaging

Confocal Z-stacks were acquired on a A1R-MP+ microscope (Nikon) equipped with 10X (air; NA 0.3); 16X (water immersion; NA 0.8) and 40X (oil; NA 1.3) objectives. Images captured in resonant scanning mode were post-processed using the denoise.ai function in NIS-Elements (Nikon). A TCS SP5 II (Leica) confocal microscope was used to capture the images in Fig. 1F,G,I,J.

### Image processing/analysis

Images in Figs 1-6 are maximum intensity Z-stack projections unless otherwise noted in the figure legends. Supplementary Movies 1 and 4 were prepared using Imaris (Bitplane) and Shotcut (https://shotcut.org/).

### Quantitative vessel analysis

Z-projections of *Tg(fli1a:EGFP)* confocal images were cropped perpendicular to the hypural diastema before being binarized with the Vessel Analysis plugin for Fiji. An individual blinded to the stage/treatment manually corrected the binary images using GIMP to accurately reflect the vessel morphology. Corrected images were input into AngioTool (Zudaire et al., 2011) to perform quantitative vessel analysis. Cropped images were 176 μm x 212 μm (width x height) in Fig. 2 and 135 μm x 426 μm (width x height) in Fig. 6.

### Quantification of innervated bony rays

Z-projections of composite Tg(sp7:mCherry-NTR), acetylated tubulin images were used to measure the number of innervated bony rays in a given fin. A bony ray was considered “innervated” if an axon traveled ≥ 50 μm through the intraray space. Z-stacks of the composite images were used to confirm that the axons were travelling in between the hemirays, not above the rays in the epidermis.

### Quantification of axon density

Z-projections of acetylated tubulin images were cropped perpendicular to hypural diastema and binarized with the IsoData threshold preset in Fiji. Axon density was quantified from these binary images using the percent area measurement in Fiji. Cropped images were 255 μm x 605 μm (width x height) in Fig. 5 and 213 μm x 605 μm (width x height) in Fig. 6.

### Statistical analysis

Scatter plots and box plots were generated with the R (https://www.r-project.org/) package ggplot2. Linear regression lines and R squared values were computed in R (ggplot, method=lm). The local polynomial regression fit in Fig. 4J was generated using the loess method. Wilcoxon rank sum test was performed in R using the wilcox.test function. Analysis of covariance (ANCOVA) between groups was computed in R using the aov function. In instances where the input data violated the homogeneity of regression slopes assumption of ANCOVA, the Johnson Neyman technique was used (R function sim_slopes).

## Acknowledgements

We thank the LSB Aquatics staff for animal care; Wai Pang Chan for imaging support; the lab of Horacio de la Iglesia for cryostat training and use; the lab of Sharlene Santana for access to computer hardware; the labs of Ajay Dhaka, Ron Kwon, Dave Raible, and Alvaro Sagasti for sharing zebrafish stocks; and Ron Kwon for comments on an earlier version of the manuscript. The authors are grateful to all members of the Rasmussen lab for discussion, technical assistance, and support.

